# CellDeathPred: A Deep Learning framework for Ferroptosis and Apoptosis prediction based on cell painting

**DOI:** 10.1101/2023.03.14.532633

**Authors:** Kenji Schorpp, Alaa Bessadok, Aidin Biibosunov, Ina Rothenaigner, Stefanie Strasser, Tingying Peng, Kamyar Hadian

## Abstract

Cell death, such as apoptosis and ferroptosis, play essential roles in the process of development, homeostasis, and pathogenesis of acute and chronic diseases. The increasing number of studies investigating cell death types in various diseases, particularly cancer and degenerative diseases, has raised hopes for their modulation in disease therapies. However, identifying the presence of a particular cell death type is not an obvious task, as it requires computationally intensive work and costly experimental assays. To address this challenge, we present CellDeathPred, a novel deep learning framework that uses high-content-imaging based on cell painting to distinguish cells undergoing ferroptosis or apoptosis from healthy cells. In particular, we incorporate a deep neural network that effectively embeds microscopic images into a representative and discriminative latent space, classifies the learned embedding into cell death modalities and optimizes the whole learning using the supervised contrastive loss function. We assessed the efficacy of the proposed framework using cell painting microscopy datasets from human HT-1080 cells, where multiple inducers of ferroptosis and apoptosis were used to trigger cell death. Our model confidently separates ferroptotic and apoptotic cells from healthy controls, with an averaged accuracy of 95% on non-confocal datasets, supporting the capacity of the CellDeathPred framework for cell death discovery.

## Introduction

Cell death can be mediated by multiple signaling pathways and each type of cell death is associated with specific changes in cell and organelle shape and cytoskeletal organization, resulting in specific morphological features. Apoptosis is the most extensively studied form of regulated cell death, but in the past two decades, other forms have been discovered, including necroptosis, pyroptosis, and ferroptosis (1, 2, 3). Cells that undergo apoptosis show these typical morphological changes: cell shrinkage followed by condensation/blistering, fragmentation and formation of apoptotic bodies (4). In contrast, other forms of cell death (necroptosis, pyroptosis, or ferroptosis) are not modulated by the activity of caspase-3/7 and hence represent distinct morphological features (5). Ferroptosis is an iron-dependent form of cell death that occurs as a consequence of lipid peroxidation (6, 7). It has been shown to be involved in multiple physiological and pathological processes, such as neurodegenerative disease, tissue damage, and acute renal failure (7). Ferroptotic cells typically show smaller mitochondria with reduced cristae and a ruptured outer membrane, but lack characteristic features of apoptosis such as chromatin condensation or apoptotic bodies (8, 9). In order to determine the acute, subacute, and chronic effects of drugs and chemical toxins, it is important to understand how a compound can induce cytotoxicity in cells and which of the various cell death pathways is activated. Cytotoxicity profiling of small molecule libraries is a well-established process in high-throughput-screening (HTS) campaigns (10), but most of the assay types detect only general cytotoxicity and do not examine the mode of action of the respective compounds. Usually, a combination of different approaches is used to study and distinguish apoptotic and the different non-apoptotic cell death processes in more detail.

It is known that cell morphology and cellular structures in response to small molecule treatment or genetic perturbations are very closely linked, such that morphological phenotypic screening might classify the mode of action of chemicals or genes in a cell (11). Based on this, the “cell painting” assay was developed. Cell painting is an image-based fluorescence microscopy assay that can be used to visualize the morphology of the cells by fluorescent labeling of cellular structures and subsequent analysis of cells (12). Through the multiplex use of fluorescent dyes, eight different cell structures can be examined simultaneously (12). After image acquisition with an automated microscope, the traditional workflow includes specialized high-content-analysis (HCA) software that can detect and further segment cellular objects and extract morphological features such as size, intensity or textures of the cell segments for further analysis including machine learning (ML) methods (12, 13). However, this requires appropriate software and can be subject to a certain bias, since only extracted features from the given images are further analyzed. Here, it is also possible that important information that would facilitate cell state classifications have not been fully detected. These features are also missing in the subsequent ML model.

Several recent publications leveraged deep learning (DL) for analyzing microscopic images and contributed a lot to canonical tasks in high-content screening (HCS) image analysis: *e*.*g*., image synthesis and feature representation (13). One example shows an image-to-image translation architecture for synthesizing three different fluorescence images from bright-field microscopy images to observe the apoptosis, nuclei, and cytoplasm of cells (14). Another study proposed a U-Net architecture to synthesize AT8-pTau image given two DAPI and YFP-tau image channels (15). With the potential of DL architectures in extracting meaningful features directly from microscopic images, recent studies proposed self-supervised learning frameworks, including a framework for studying the temporal drug effect on cancer cell images, or a framework to learn phenotypic embeddings of HCS images using self-supervised triplet network (16, 17). While these advancements in DL application to HCS images offer the potential to accelerate drug discovery, so far there is only very little work about the analysis and prediction of regulated cell death. Understanding and identifying drugs that lead to distinct cell death modalities is of high importance. Two studies describe ML and DL methods for predicting cell death modalities using microscopic images. The first work leveraged multinomial logistic regression models using the LASSO for discriminating microscopic images of fluorescently stained cells undergoing different cell death modalities–ferroptosis and apoptosis (18). Although promising, the current model is based on specific immunostaining, TfR1 (19) and Hoechst, which is limiting its generalizability to other cell death modalities. The second work utilizes a VGG-19 deep network to discriminate apoptosis from necroptosis (20). This method proposed a pre-filtering step to filter all cells that showed alive morphology from cell images where inducers were added, which enforces the DL model to classify cell death modalities from well selected image features.

In this study, we demonstrate a framework that learns from cell painting images without any pre-filtering step. This not only reduces the extra computation of a filtering step, but importantly, it enables the model to learn from heterogeneous cell images; thus, being a more generalizable model for different kinds of cell death-related images. The present work deals with the question: “Given microscopic images generated from a high-content cell painting assay, can we classify whether the drug induces ferroptosis, apoptosis or has no adverse effect?” Addressing such a question may be important in predicting the presence of a particular cell death type in clinically relevant drugs, which may open up new therapeutic possibilities.

## Results

### Characterization of ferroptosis versus apoptosis inducers

For this study, seven well-characterized apoptosis and ferroptosis inducers (FINs) were used to explore the applicability of a DL framework to classify ferroptotic, apoptotic and healthy cells (**Fig. 1A**). In order to confirm that ferroptosis is indeed induced by FINs, HT-1080 fibrosarcoma cells were seeded and either untreated or pre-treated with the ferroptosis inhibitor ferrostatin-1 (Fer-1) before RSL3 was added as a representative FIN at various concentrations (20-point titration). Fer-1 is described as an inhibitor of lipid peroxidation and rescues cells from ferroptosis (3). HT-1080 cells were chosen because they are well established in ferroptosis research (3, 21, 22) and well-suited for microscopy. By using the CellTiter-Glo (CTG) viability assay, which measures intracellular ATP levels, it could be shown that co-treatment of FINs with Fer-1 rescued cells from undergoing cell death (**Supplementary Fig. 1**). In contrast, staurosporine (STS) induced cell death could not be rescued with Fer-1 co-treatment (**Supplementary Fig. 1**), demonstrating that the selected molecules are specific.

**Figure 1:**
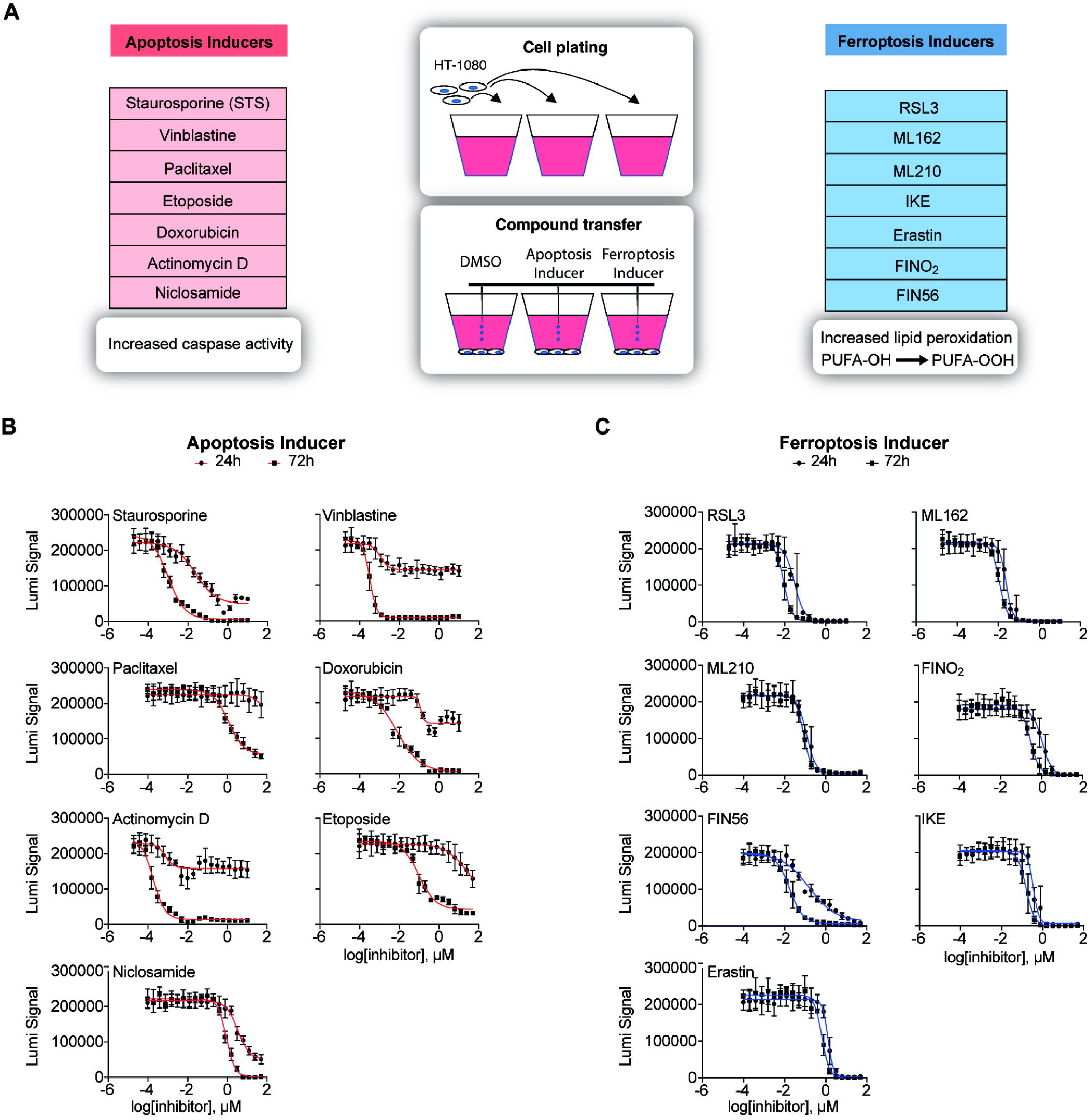
Identification of the ideal concentrations of apoptosis and ferroptosis inducers. **(A)** Schematic overview of the cell death inducers used for this study. HT-1080 cells were seeded and treated with 7 apoptosis inducers, 7 ferroptosis inducers (FINs) and DMSO as a solvent control. Cells treated with apoptosis inducers execute the apoptotic program by activating caspases. Treatment of cells with FINs result in lipid peroxide accumulation due to the limited GPX4 activity and hence induce ferroptosis **(B)** Results of the dose response (20-point) viability assay with apoptosis inducers in HT-1080 cells. 24h and 72h incubation time. Cellular ATP levels were measured using luminescence signals. Values indicate mean ± SD (n = 6, technical replicates). **(C)** Same as in (B) here for treatment with FINs.

### Experimental setup and imaging upon cell painting

For subsequent experiments, it was important to choose compound concentrations that have a mild to moderate effect on cell viability. Our goal was to treat cells in such a way that cell death was induced (ATP reduction), but the cells are not yet affected by excessive end phase necrosis. For this, we performed a pilot study with a wide range of concentrations (20-point titration) to determine optimal concentrations for each of the 14 small molecules. (**Fig. 1B**). We measured intracellular ATP levels after 24 and 72 hours, respectively, and then selected a concentration spectrum of 5 concentrations based on the IC_50_ value for the subsequent experiments (**Table 1**). Different treatment durations of 24 and 72 hours were chosen, because some substances are known to induce cell death very quickly, while other compounds only have an increased toxic effect after several rounds of the cell cycle.

**Table 1:** Five concentrations for each substance. IC50, one concentration higher [+] and three different concentrations lower [-], [- -] and [- - -] than the IC50

Guided by the results of the pilot study, cell painting experiments were conducted using the optimized compound concentrations. HT-1080 cells were seeded in 384-well plates and treated with the five defined concentrations of the cell death inducers (**Table 1**) for 24 and 72 hours. We used five (Hoechst 33342, Wheat Germ Agglutinin, Concanavalin A, TRITC-Phalloidin, Mitotracker) instead of the six dyes, which are described in the standard cell painting protocol, allowing us to run the assay on only one plate per data point instead of two parallel plates. Image sets of two slightly modified independent experiments were collected as training data sets: in experiment 1 the cells were imaged with 40x magnification and a confocal spinning disk, while for experiment 2 a 40x magnification and widefield was used. Nine technical replicate wells for each substance and concentration within one experiment were imaged. In addition, we recorded nine to eleven images per well using four different fluorescence channels, creating a large image data set per experiment. In order to check the strength of cell death for each treatment condition (compound and concentration), we performed CTG assays in parallel (**Fig. 2A**). Importantly, the cell pool, the number of cells, the compound plates, and the way of treatment that were used for CTG measurement were identical to those used in the cell painting experiment. In a next step, the data of the ATP measurement and cell paining were annotated. By this we were able to select only images for training that reflected a certain level of intracellular ATP reduction. In all the experiments, we normalized the absolute ATP values to the DMSO levels. So, the wells with values close to 1.0 are considered not to be affected. While those approaching 0.0 are “dead”. We selected wells with ATP values that fall into the range [0.3 - 0.8]. Supposedly, in these wells the treatments did not completely kill the cell population and at the same time caused some non-negligible effect including morphological changes. We also analyzed the images with the Columbus high-content-analysis software. For this purpose, the nuclei were identified using Hoechst signal, and based on this, the cytoplasm and the membrane regions were segmented using the F-actin signal (**Supplementary Fig. 2**). The intensity, the morphology, and the symmetry of the objects, as well as the texture properties, and structure of the fluorescence signal were determined within these defined cell segments for the different fluorescent channels. This resulted in 245 extracted features that could be used for classical machine learning (ML) approaches. Importantly, the features for single cells were averaged for images coming from the same well (median).

**Figure 2:**
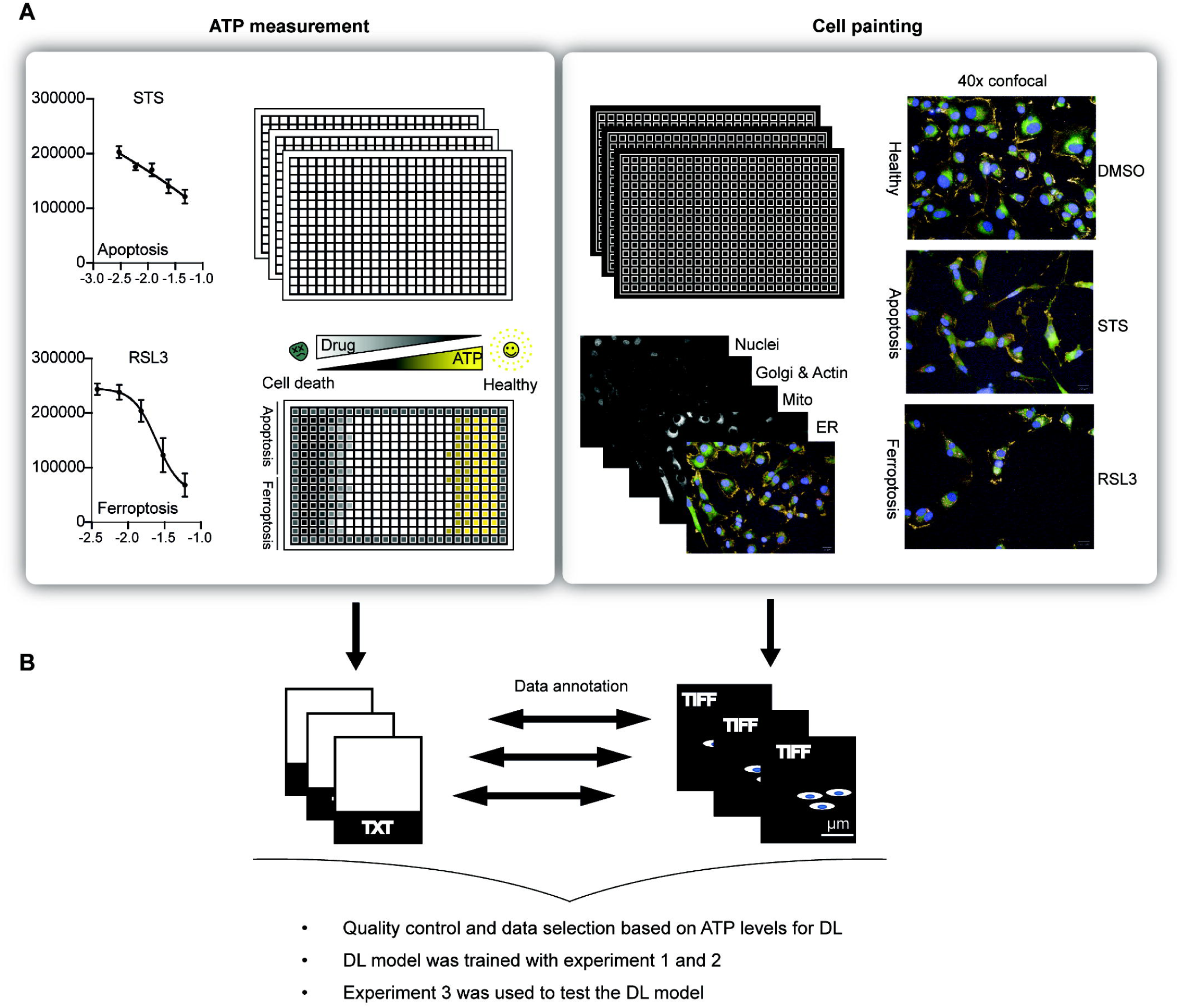
Schematic overview of the data generation process. **(A)** HT-1080 cells were treated with five different concentrations of apoptosis and ferroptosis inducers. ATP measurement (left) and cell painting (right) experiments were conducted in parallel. Staurosporine (STS) and RSL3 data are shown as representative data for apoptosis and ferroptosis inducers, respectively. Values indicate mean values ± SD of 6 technical replicates. The cells were imaged with a 40x objective. The different organelles (nuclei, golgi apparatus, actin cytoskeleton, mitochondria, endoplasmatic reticulum) were imaged using four different fluorescence channels. **(B)** The data from the viability assay were annotated with the images from the cell painting experiment. Only if viability was in the range of 80-30% the images were used for model training. Three experiments were performed. Experiments 1 and 2 were used for training the CellDeathPred model. Experiment 3 to test the model.

### CellDeathPred architecture and classification strategy

To predict whether treatment of cells with a certain drug induces apoptosis or ferroptosis or has no effect, we developed CellDeathPred, a DL architecture comprising four parts: data augmentation, model backbone, an embedder network trained with supervised contrastive loss and finally a classification network trained with cross-entropy loss (**Fig. 3A**). Data augmentation is a widely used technique in DL, which aims to improve the generalizability of the model during training; thus, enhancing the prediction accuracy of the classifier. We applied five crops to the initial 1320×1024 image. Then, we applied to each 512×512 crop the horizontal and vertical flips, 90 rotation and the gaussian noise augmentations (**Fig. 3B**). We chose our model backbone to be EfficientNet-b0 (23), a convolutional neural network that is pretrained on the large scale of the ImageNet dataset comprising 1,000 classes of RGB images. Since our images have four channels (*i*.*e*., ER, Actin/Golgi, Mitochondria, and Nuclei), we replaced the first layer of the backbone network, which originally accepts three-channel images with a four-channel input. We assigned the average weight of the three channels to be the weight for the fourth channel. The pretrained EfficientNet-b0 has a role of feature extractor for our cellular dataset; therefore to classify the drugs we added a classification layer trained with the cross-entropy loss. While this is a universal loss term used in most of DL classification frameworks, recent studies showed that the cross-entropy loss alone cannot guarantee a good generalizability of the trained network, in particular in the presence of a batch effect (24). Batch effect is a common problem in microscopy imaging data, which refers to systematic differences such as temperature or microscopy lighting conditions in an experiment cause change in the image intensities and features from batch to batch–*i*.*e*., one batch refers to a set of experimental plates that are executed together. To solve this issue, we added a supervised contrastive loss (SupConLoss) after an embedder network that consists of a sequence of fully connected layers that maps the output of the backbone into a low dimensional space (Fig 3A). SupConLoss is a recent state-of-the-art metric learning loss term that aims to maximize the similarity between a pair of samples in the same class whilst minimizing the similarity of two samples from different classes (25). By encouraging the network to learn a more robust and discriminative representation, SupConLoss improves its generalizability (for more details we refer the reader to the material and methods section). Moreover, to better overcome the batch effect issue, we further propose a batch-aware sampling strategy in conjunction with SupConLoss (for more details we refer the reader to the material and methods section). A byproduct of SupConLoss is image retrieval at the testing stage (**Fig. 3C**): given an input testing image our CellDeathPred framework generates the embedding features and retrieves K-nearest-neighbor (KNN) in the training images by ranking feature distance in the embedding space (we choose $k = 1$ and a cosine distance as a metric, details refer to the material and methods section). Besides image retrieval, CellDeathPred also predicts the probability of an input image to belong to each of the main classes (i.e., healthy, apoptosis or ferroptosis) (**Fig. 3D**).

**Figure 3:**
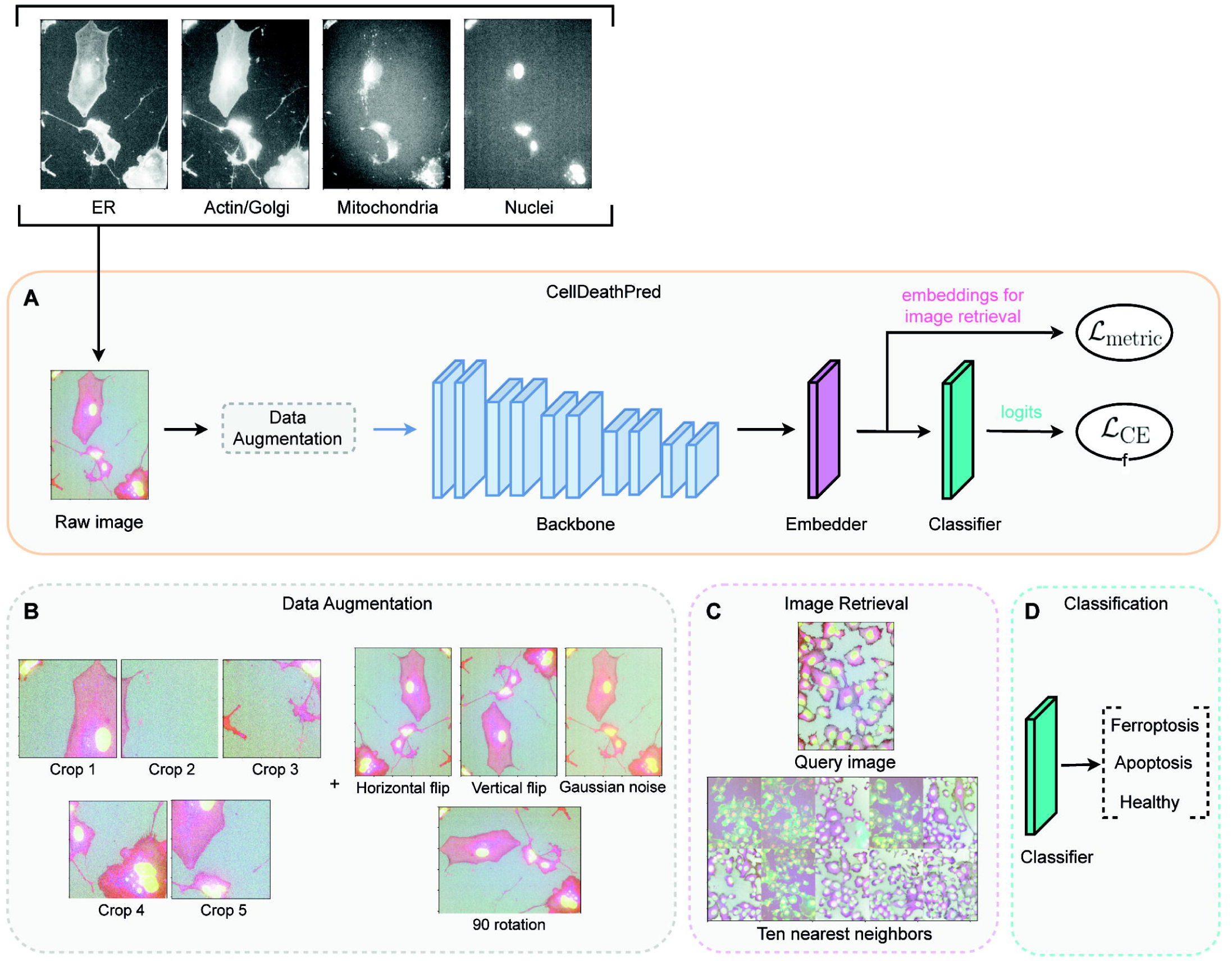
CellDeathPred architecture for classifying cell death modalities. Given four channels of the raw image (ER, Actin/Golgi, Mitochondria and Nuclei) as input, the neural network predicts whether the drug used in the experiment induces apoptosis or ferroptosis or it is a DMSO one. The architecture comprised four phases: data augmentation to ensure robustness of the model during training, (2) backbone model which is a pretrained network (Efficientnet-b0), (3) an embedder is a sequence of fully connected layers to map the input data to a low-dimensional space, and (4) a classifier which is a sequence of fully connected layers that outputs the predicted modality. **(B)** Illustration of the used augmentations. Four corner and one center crops with sizes 512×512. Augmentations were applied to each crop. **(C)** Example of an image retrieval. Ten nearest neighbors for a query image in the embedding space. **(D)** The last layer with three nodes of the model. Classification predictions of the three classes.

### DL versus ML classification to classify cell death from cell painting

In order to test the CellDeathPred model on a previously unseen data set, we performed a third experiment in the same way as described in Figure 3 (**Supplementary Fig. 4**). In contrast to experiment 1 and 2, the cells were only treated for 24 hours, but this time we imaged the plates of the same experiment both confocal and non-confocal to investigate whether this has an impact on the classification accuracy. Also, these images were analyzed with our imaging software and 245 features were extracted. Before evaluating ML models on the features extracted by Columbus software, a preprocessing step was essential for training the models. Mainly, we removed columns with not a number (NaN) values, duplicated columns and normalized the values. We chose three ML models widely used in the literature: Random Forest (26), Logistic Regression (27) and AdaBoost (28). We used uniform manifold approximation and projection (UMAP) (29) as a dimension reduction technique to visualize how images of ferroptosis drugs cluster from those of apoptosis, and the healthy cells. As shown in the UMAP of CellDeathPred learnt feature space (**Fig. 4A (DL)**, images of healthy, apoptosis and ferroptosis classes are clustered into three distinct clusters, whilst they are mixed in the UMAP of cellular features extracted from Columbus software (**Fig. 4A (ML)**). A side-by-side comparison of DL vs ML methods (**Fig. 4B**) show that CellDeathPred reached a classification accuracy of over 93% for Plate 1 and Plate 2, which is almost 10% increase over the best ML method. Also, in most cases, CellDeathPred with SupConLoss outperforms CellDeathPred w/o SupConLoss (**Fig. 4B**), suggesting better model generalizability. This could also be demonstrated by the UMAP of feature space when we color the images according to their batch whilst images of different plates are well mixed, suggesting SupConLoss is effective in avoiding the batch effect problem (**Supplementary Fig. 5**).

**Figure 4:**
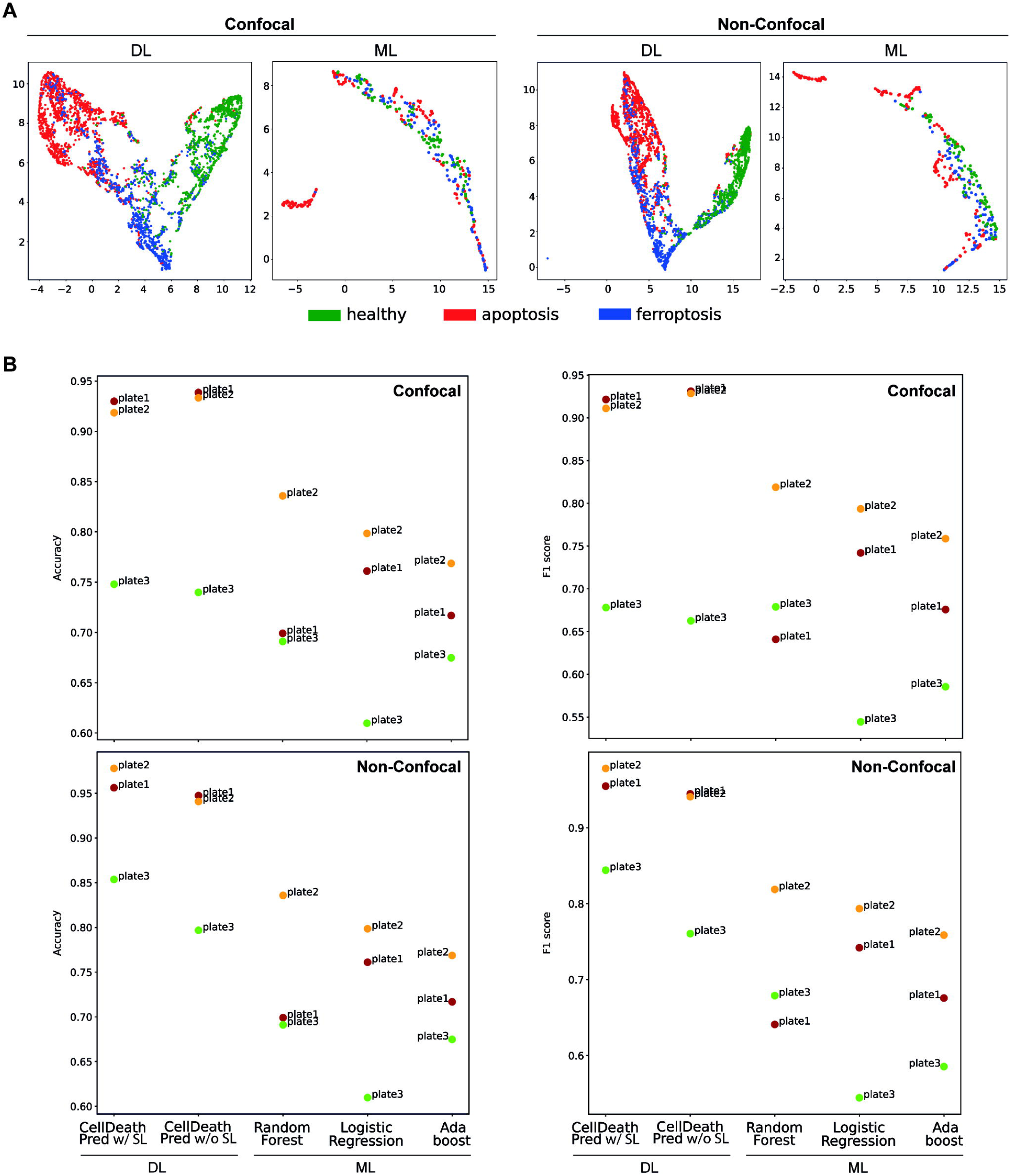
Comparing CellDeathPred with other ML models. **(A)** UMAP of embeddings of experiment 3 plates with confocal and non-confocal imaging. Every point corresponds to the embedding of an image. On the left using the CellDeathPred model which was trained on images from experiment 1 and 2. Individual wells are visualized as points on the scatter plot of the first two principal components. On the right UMAP of 245 features extracted from the images initially extracted from Columbus software. The color code is according to the drug category (blue = “healthy”, red = “apoptosis”, green = “ferroptosis”) and was added after the UMAP was conducted. **(B)** Accuracy and F1-score results on the well level are shown per plate for confocal data (top row) and non-confocal data (on the bottom row). The x-axis represents the proposed method CellDeathPred, its variant where we remove the SupConLoss and the machine learning-based methods widely used in the literature. The proposed deep learning model achieved best performance in both evaluation metrics compared to the comparison methods.

### Classification of ferroptotic and apoptotic cells at different drug concentration using CellDeathPred

In the previous paragraph, we demonstrated that CellDeathPred is more accurate than the tested ML models. Moreover, we have shown that non-confocal images are sufficient for the separation of the different classes in the UMAP and for accurate classification. Therefore, we focused on the DL results of the non-confocal images as their acquisition requires a fraction of the imaging time compared to confocal images with comparable prediction accuracy (**Fig. 4B**). Like all other experiments, experiment 3 was performed as technical triplicate (three different 384 well plates, 24h treatment). In addition, every substance, whether ferroptosis or apoptosis inducer, occurred in triplicates in five different concentrations on each of the three plates. First, we checked the ATP level from the experiment that was conducted in parallel to the cell painting assay. Most of the 14 substances led to increased cell death with increasing concentrations (**Supplementary Fig. 3**), indicating that we have chosen the correct concentration range. The confusion matrix of the CellDeathPred DL model showed that the prediction worked very well, and to a large extent both types of cell death inducers and healthy cells were correctly classified (close to 100% for plates 01 and 02). However, it also showed that the predictions for plate 03 were worse than for plate 01 and 02 (**Fig. 5A**). To identify the cause of the lower prediction accuracy of plate 03, we used the data from the image segmentation and the feature nucleus area as an indicator of the potency of the small molecules in the cell painting experiment (**Fig. 5B**). Here, plate 03 appeared to be very different from plate 01 and 02, where cell death induction is much weaker. In fact, by chance plate 03 acted as a good control and confirmed the high accuracy of CellDeathPred that classifies cell death only if sufficient loss of viability is present, which was not the case in plate 03. Next, we analyzed the prediction accuracy of the individual substances depending on the ATP signal. Usually, a correct prediction was achieved with a cell viability of around 50% (**Fig. 5C**). If the concentrations of the substances are too low and thus the viability higher than 80%, the cells are classified as healthy. For example, the ATP levels in cells treated with Actinomycin D and Erastin are relatively high in this experiment, indicating that they did not induce cell death (**Fig. 5C**). Accordingly, CellDeathPred classified the cells treated with Erastin or Actinomycin D mainly as “healthy” in this case, again demonstrating its high accuracy based on the experimental perofrmance. Together, we have developed CellDeathPred, a DL framework, which is able to classify ferroptotic and apoptotic cells with an accuracy of close to 100% using non-confocal images, when the drugs sufficiently induce the type of cell death.

**Figure 5:**
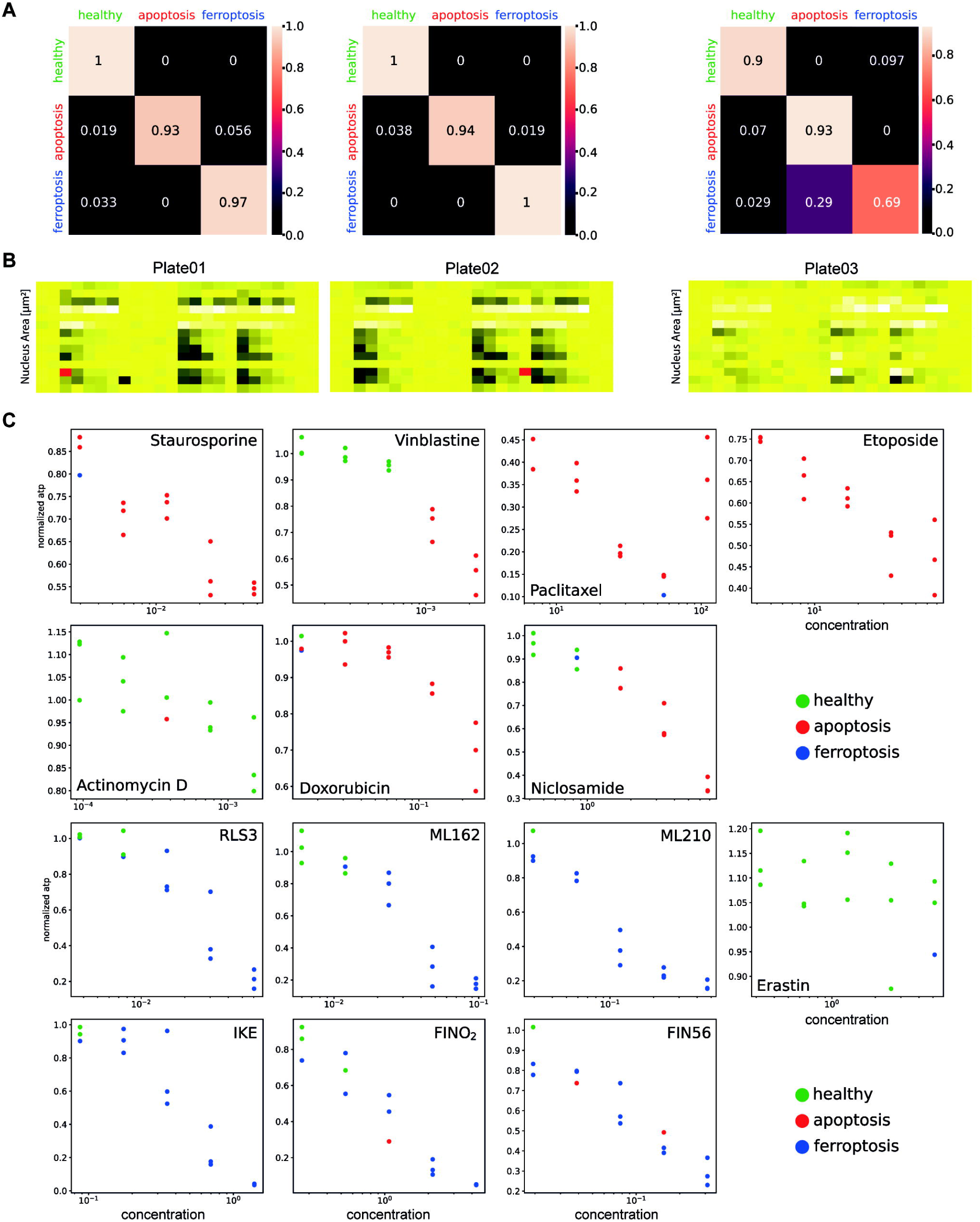
Results of the DL model. **(A)** Confusion matrices for experiment 3 (non-confocal) with CellDeathPred model. The model was trained on images from experiment 1 and 2. Order of plates from left to right. Heatmap of the ATP measurement that was conducted in parallel to the cell painting experiment. Low (black) and high (yellow) luminescence signals correspond to the cellular ATP levels. The experiment was performed in technical replicates (three plates). Cells were treated with five different concentrations for each small molecule. **(B)** Heatmap of the nuclei count that was conducted on the images of the cell painting experiment. Low (black) and high (yellow) luminescence signals correspond to the number of selected nuclei. The experiment was performed in technical replicates (three plates). Cells were treated with five different concentrations for each small molecule. (**C) (C)** Prediction of every substance for every concentration across the plate depending on the ATP level (normalized). Performed for experiment 3, plate01 (non-confocal), with the CellDeathPred model. For every concentration there are three replicates.

## Discussion

To induce cell death, we chose seven apoptosis inducers and seven FINs to ensure that the generated data are balanced for each of the respective cell death modality. Notably, we selected the cell death inducers to modulate different biological targets of the given cell death modality in order to cover larger aspects of apoptosis and ferroptosis. Here, the seven apoptosis inducers targeted caspases, microtubules, oxidative phosphorylation, RNA-synthesis, and topoisomerase II. The seven ferroptosis inhibitors belonged to class I, II, III and IV FINs; thus, inhibiting system x_c_^−^, and GPX4 activity in a direct or indirect manner. Of course, the collection of 14 small molecules does not cover all possibilities to induce apoptotic or ferroptotic cell death, and it may be interesting to test other substances in future to understand if CellDeathPred would correctly classify them into the correct categories.

We created the CellDeathPred model by using datasets in a single cell line and one type of microscope with specific settings, and we are aware that this choice limits the potential for generalizability of our model using it for datasets created with other cell lines and microscopic devices. However, the 14 selected substances can serve as internal benchmarks to generate comparable datasets with other cell lines and other small molecules that can be used to train the model.

Determining the exact concentration series of the different substances was crucial to generate highly standardized data in order to minimize technical variability and therefore maximize the biological signal in the data. Our assumption was that if the concentrations are too low the cells are relatively healthy. In contrast, if the concentrations are too high the cells might be already in a necrotic phase and any kind of ML or DL model would have problems to correctly classify these cells. In fact, we recommend identifying the IC_50_ for all substances used in new experiments in order to have internal controls for the assay. With the defined induction rate, we made use of the cell painting assay (12) to visualize healthy, apoptotic or ferroptotic cells. Previous efforts to stain cell death have been selective of a given cell death modality, *e*.*g*., *TfR1* staining for ferroptosis (18, 19) or Annexin-V staining for apoptosis (30). Importantly, by applying cell painting to visualize cell death the procedure does not rely on specific markers, but can use general content-information about DNA, ER, mitochondria, Golgi, and actin to profile cells with regards to distinguishing healthy state from apoptosis and ferroptosis. This advance will enable the rapid transfer of CellDeathPred to other forms of cell death, such as necroptosis and pyroptosis. The performance of CellDeathPred to classify apoptosis and ferroptosis was close to 100%, and importantly in cases where experimental failure led to poor cell death induction (*e*.*g*., Erastin hardly induced any ferroptosis in plate 03 of the test experiment) the model correctly classified such samples as healthy cells (**Fig. 5C**).

In CellDeathPred, we applied contrastive learning to allow the model to pull together cell death modality images of a particular treatment. This is represented in the total loss that combines the contrastive learning and cross-entropy losses with equal weights. Moreover, to combat the batch effect during training, we added more diversity by including images from different plates to a batch and thereby reducing batch effect. Our choice for the backbone model was also important to achieve high prediction performance. In the literature, EfficientNet models demonstrated a better efficiency over existing state-of-the-art architecture such as ResNet-18 (31). Therefore, adopting a transfer learning strategy and further training the backbone on our cellular dataset had the advantage regarding computational efficiency and accuracy compared to training from scratch and extracting representative features. As our model is the first deep learning model aiming to distinguish ferroptosis-based modulators from apoptosis and healthy ones, we compared it with a baseline method “CellDeathPred w/o SL”, where we removed the SupConLoss and we kept the same architecture, and three different machine learning models Random Forest, Logistic Regression, and AdaBoost. To validate our model, we investigated the accuracy and F1-score metrics for the four comparison methods (**Fig. 4**). Our model (“CellDeathPred with SL”) mostly outperformed its variant (w/o SL), and was better than all ML models in confocal and non-confocal assays.

It would be interesting for the future investigations to investigate substances that have not yet been assigned to any of the cell deaths, but which have shown cytotoxic effects, and understand if they induce a specific form of cell death. Moreover, CellDeathPred should be expanded in future studies to integrate other regulated cell deaths, such as necroptosis and pyroptosis.

## Conclusions

In summary, we have demonstrated that our CellDeathPred framework is able to accurately classify cells that were treated with small molecules inducing ferroptotic and apoptotic cell death. Here we present a detailed experimental protocol on how to generate the data, to train and use our developed CellDeathPred model. This work will contribute to the characterization of cell death inducing small molecules or biologics, and thereby help to better understand their mode of action. We think that our work based on the cell painting protocol in combination with our DL model can be extended to other questions in the classification of chemical substances and thus may act as a blueprint for comparable future projects.

## Materials and methods

### Cultivation of cells

HT-1080 cells were cultivated in DMEM with high glucose, glutamine, and pyruvate (Gibco™), supplemented with 10% FBS (Gibco™), 1% Penicillin-Streptomycin (Gibco™), 1% NEAA (Gibco™). They were grown in the incubator at 37 °C and 5% CO_2_.

### CellTiterGlo® assay

HT-1080 cells were seeded 1000 cells/well in 50 µl DMEM with high glucose, glutamine and pyruvate (Gibco™), supplemented with 10% FBS (Gibco™), 1% Penicillin-Streptomycin (Gibco™), 1% NEAA (Gibco™) on white opaque 384-well CulturPlate-384 Microplates (PerkinElmer, 6007680). Seeding was performed with the MultiFlo Microplate Dispenser (BioTek). The next day compounds were diluted in DMSO on a compound plate. 0,5 µl were transferred from the compound plate onto the cells with the Sciclone G3 Liquid Handling Workstation (PerkinElmer). Before addition of compounds, cells were pre-treated with 5 µl media (control) or 5µl Fer-1 media solution to have a final concentration of 2 µM on the cells. When the CellTiter Glo assay was performed in parallel with the cell painting assay, deviations from the standard assay protocol occurred. 1000 cells were seeded in 25 µl media instead of 50µl. In addition, as for the cell painting experiment, an intermediate dilution step was introduced during compound transfer. For this, compounds were transferred into plates containing only cell medium. In a next step 25 µl of the compound cell culture media mix were carefully transferred from the intermediate plate on plates with the cells in 25 µl media using a Beckman Coulter Biomek Fx. After 24 h or 72 h incubation at 37 °C and 5% CO_2_ in the incubator (Cytomat, ThermoFisher), 25 µl CellTiterGlo® (Promega) per well was added and the luminescence was read at 700 nm, measurement height 6.5 mm and measurement time 0.5 s with EnVision Multimode Plate Reader (PerkinElmer).

### Cell painting Reagents

Mitotracker Deep Red (Invitrogen, #M22426), WGA (Invitrogen, #W32464), Concanavalin (Invitrogen, #C11252) and Hoechst 33342 (Invitrogen, #H3570) stock solutions were prepared according to supplier information. For Phalloidin-TRITC (Sigma, #P1951) methanol was added to 1 vial to prepare 0,1 mg/ml stock solution.

### Cell painting assay

HT-1080 cells were seeded with a cell number of 1000 cells/well in 25µl cell culture media on PhenoPlate^™^ 384-well microplates (PerkinElmer, 6057308) with the MultiFlo Microplate Dispenser (BioTek) to have 6 replicates. Compounds were diluted in a 20- or 5-point titration with DMSO (x mM) as highest concentrations on a compound plate. On the next day transfer and mix of compounds into plates only containing cell culture media was performed with the Sciclone G3 Liquid Handling Workstation (PerkinElmer). This intermediate dilution step was included to avoid DMSO gradient effects on cell monolayers. In a next step 25µl of the compound cell culture media mix were carefully transferred from the intermediate plate on plates with the cells using a Beckman Coulter Biomek Fx followed by incubation in the Cytomat incubator (ThermoFisher) at 37°C and 5% CO_2_ for 24 and or 72 h. The cell painting protocol was performed as described in Anne Carpenter’s original publication (12). The only deviations were the omission of the Syto14 dye and the use of phalloidin-TRITC instead of phalloidin-568. The settings we have chosen for the Operetta microscope are as follows: Acquisition type: Spinning disk confocal or widefield with 40x high na objective. Main emission [nm]/main excitation [nm]: 525/475 for ER stain, 445/380 for nucleus stain, 705/630 for mitochondria stain, 595/535 for actin-RNA stain.

### Automated image analysis

Image analysis was performed using Columbus software version 2.9.1 (PerkinElmer). In the following, the analysis steps in Columbus are described: the Hoechst 33342 and TRITC signals were smoothened for the cell segmentation process using Median filters to reduce noise signals. Nuclei were detected via the Hoechst 33342 signal. The TRITC channel was used to define the cytoplasm. In a next step, morphology/symmetry features, texture (SER features), and intensity properties of the Hoechst 33342, TRITC, 488 and 647 channels were calculated for each cell region (nuclei and cytoplasm). Moreover, we applied a filter to remove border objects (nuclei that cross image borders). For the detailed analysis pipeline in Columbus, please see **Supplementary Table 1** with the analysis sequences.

### Model training and application

Designing a DL model for distinguishing between cell death modalities given a set of microscopic images is challenging due to several factors as explained previously: batch effect and reduced generalizability performance. To overcome these issues, we propose to train in a supervised contrastive learning fashion using the SupConLoss defined as follows:

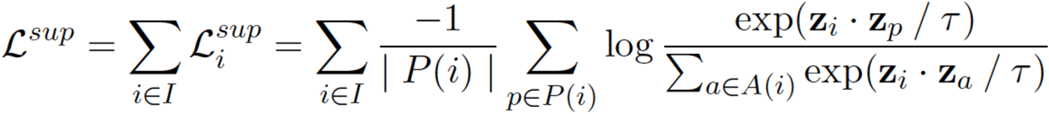

$z_i$ is an embedding vector with class label $y_i$ generated by the embedder network. $i$ is an anchor in the batch $I$. $P(i) = {p ∈ A(i) : y_i = y_p}$ denotes positive samples that belong to the same class. SupConLoss first calculates the inner product of the anchor with samples in $P(i)$, second applies an exponential function in order to amplify large values. The outputs are summed up and normalized over all samples $A = I\(1)$. $τ$ denotes the supervised temperature, a hyperparameter that helps disentangling positive and negative samples. The main benefit of our supervised contrastive learning is that it disentangles batch effects from relevant biological variables (24, 32) and this can be seen in the UMAP reported results (**Fig.4 A**). Mainly, we chose UMAP over other dimensionality reduction techniques such as t-distributed stochastic neighbor embedding (t-SNE) or principal component analysis (PCA) owing to its speed and performance for the preservation of the global structure of the data. To further tackle the batch effect problem, we propose a batch-aware sampling strategy to better train our network. We further use categorical cross entropy as classification loss during training. Ultimately, in our CellDeathPred architecture, we define the overall loss function as:

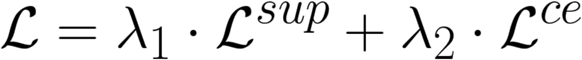

where $\lambda_1$ and $\lambda_2$ are hyper-parameters that control the relative importance of SupConLoss and cross entropy losses, respectively. Empirically, we set the temperature parameter of the SupConLoss, the learning rate and both $\lambda_1$ and $\lambda_2$ to 0.1, 1.25e-5 and 0.5, respectively. We define the batch size $bs$ using the following formula:

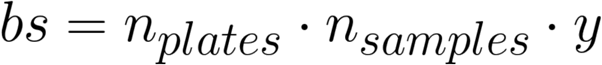

For a batch of size 30, we compute it as 30 = 2 * 5 * 3, which means it consists of samples from two plates with 5 samples of each cell death modality. The batch-norm layers were freezed to reduce overfitting. In the contrastive learning context we define a sample triplets: anchor, positive and negative. We define the anchor as an image belonging to a plate $p_i$ and the class $y_i$, while we choose the positive sample to be an image from another plate $p_p$ ($p_i **≠** p_p$) having the same label as the anchor and we define a negative sample as an image belonging to the same plate of the anchor with a different label $y_n$ ($y_i **≠** y_n$). The ultimate goal is to minimize the distance between the anchor and positive samples and maximize the one between the anchor and negative samples. The main advantage of the sampling strategy is increasing heterogeneity of the batch during training thus increasing the generalizability of the model in the case of having a dataset with batch effect. To evaluate our model, we first train it using 80% of the dataset and test it on 20%. We report accuracy and F1-score results in the field level (**Fig. 4B**) and in the well level (**Fig. 5A**). Knowing that all fields belonging to a particular well have the same label, we define the prediction on the well level as the majority voting where we count the number of apoptosis, ferroptosis, healthy predictions of the fields and assign the class with the maximum votes as the well class.

Equipped with the above components, the proposed CellDeathPred not only overcomes the issues of applying a DL model on cell painting data but also represents the first automatic labeling method of drugs that can be easily adopted for classification on other datasets.

## Acknowledgements

We thank Christian Pütz and Stefanie Brandner for excellent technical assistance.

## Author Contribution Statement

K.S. and K.H. conceptualized the study; K.S., I.R., S.S., and K.H. performed and analyzed the wet-lab experiments (viability assays, cell painting, imaging, feature extraction for machine learning); A.B., A.B., and T.P. generated the CellDeathPred deep learning framework, analyzed the cell paining data (deep learning and machine learning), and classified cell death modalities. K.S., A.B. (Alaa), T.P., K.H. wrote the original draft; All authors read and edited the manuscript, commented and approved the manuscript for submission.

## Ethics Statement

No ethical concerns.

## Data Availability Statement

The image data used in this work are available at https://github.com/peng-lab/CellDeathPred/tree/main/Dataset and the source code for the models used in this work is available at https://github.com/peng-lab/CellDeathPred/tree/main/Code.

