## Supplementary Figures for "CellDeathPred: A Deep Learning framework for Ferroptosis and Apoptosis prediction based on cell painting"

### SUPPLEMENTARY FIGURE 1

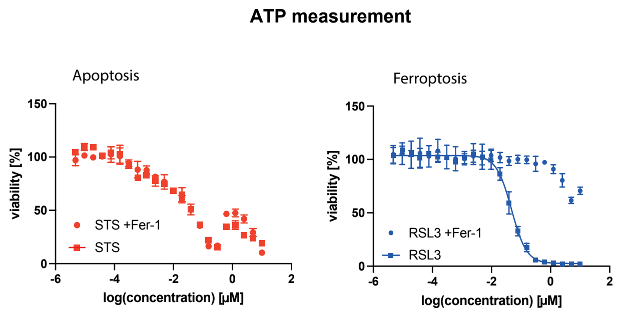


**Supplementary Figure 1: RSL3 and not STS induce Ferroptosis in HT-1080 cells**.
Dose response (20-point) viability assays with RSL3 as representative for FINs and STS as representative for apoptosis inducers with or without Fer-1. 24 h incubation time. Cellular ATP levels were measured using luminescence signals. Values indicate mean ± SD (n = 3).

### SUPPLEMENTARY FIGURE 2

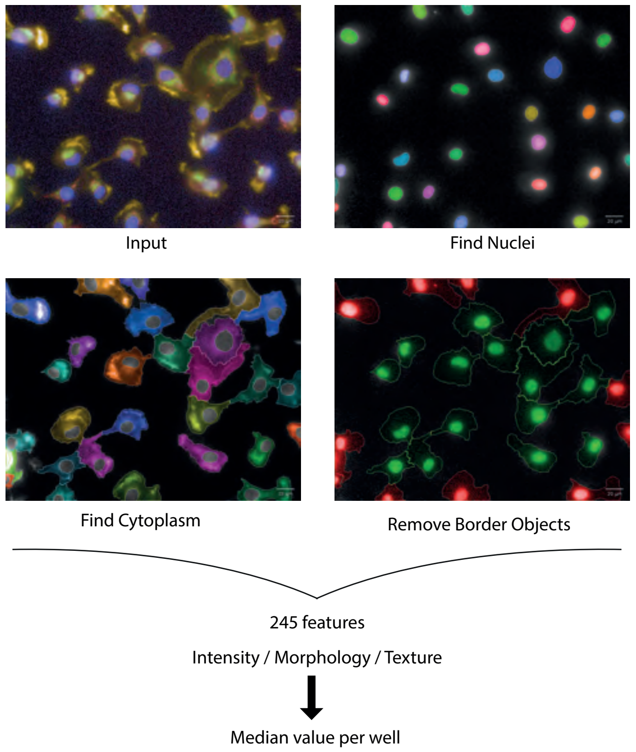


**Supplementary Figure 2: Schematic overview of the different steps of the Columbus analysis pipeline.**
The cells on the input images were segmented into Nuclei and Cytoplasm.
Border objects were removed and then features for intensity, morphology and texture were extracted.

### SUPPLEMENTARY FIGURE 3

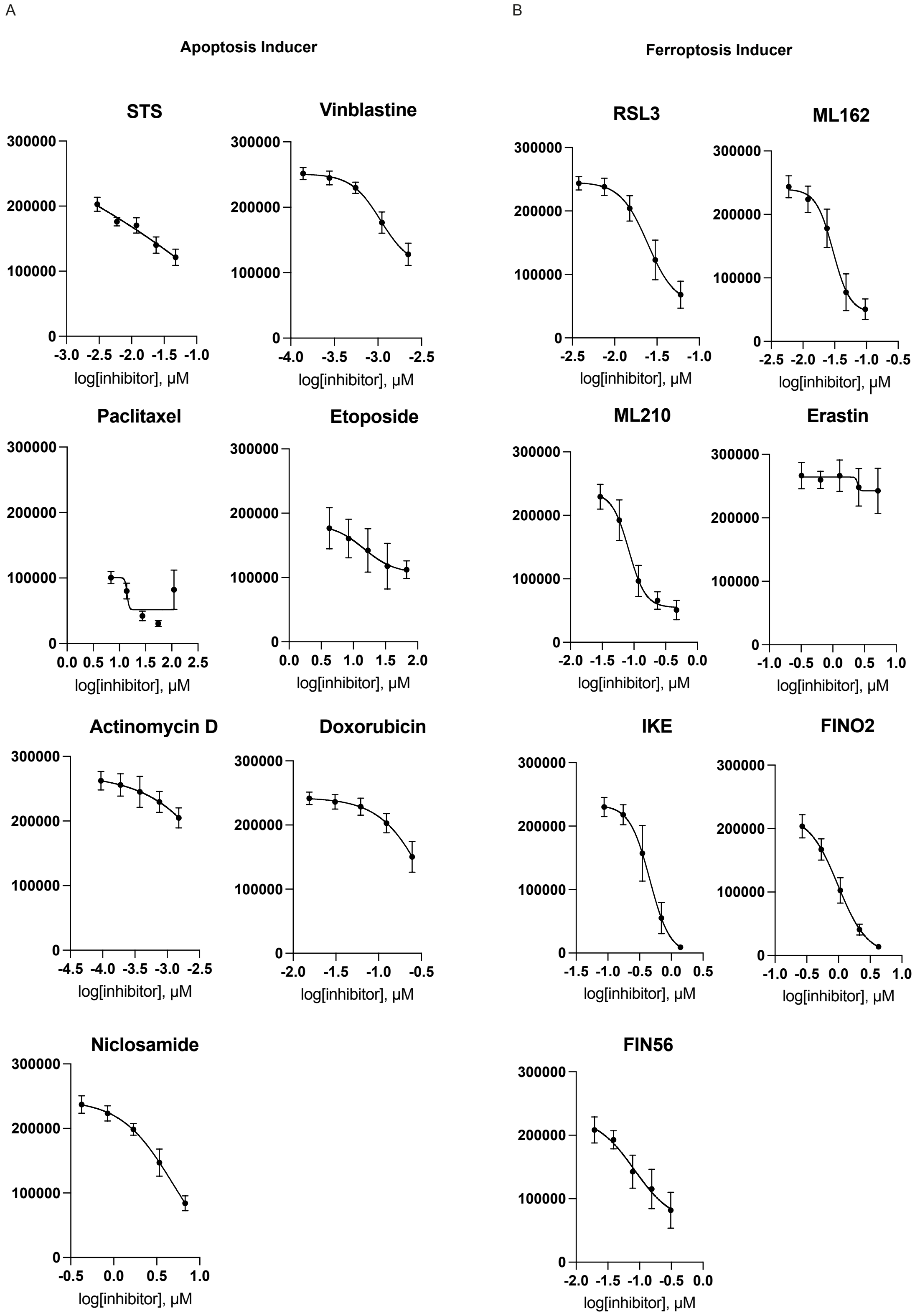


**Supplementary Figure 3: Results from the cell viability assay for experiment 3 after 24h incubation with treatments in HT-1080 cells.**

Values indicate mean ± SD (n = 6).

### SUPPLEMENTARY FIGURE 4

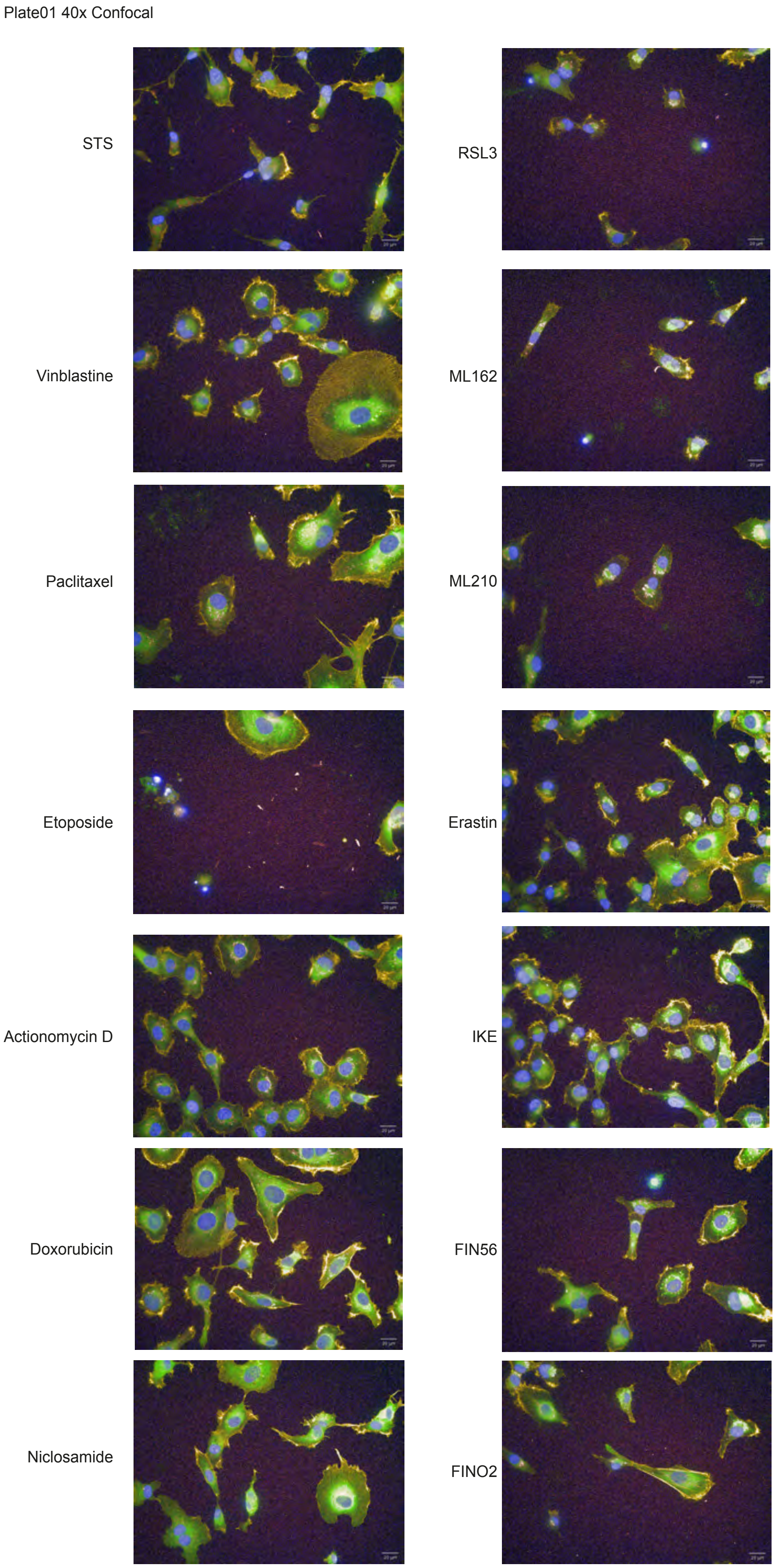


**Supplementary Figure 4: Representative images of the cell painting experiment 3.**
(Plate01, 40x objective, confocal) for all the different treatment conditions.

### SUPPLEMENTARY FIGURE 5

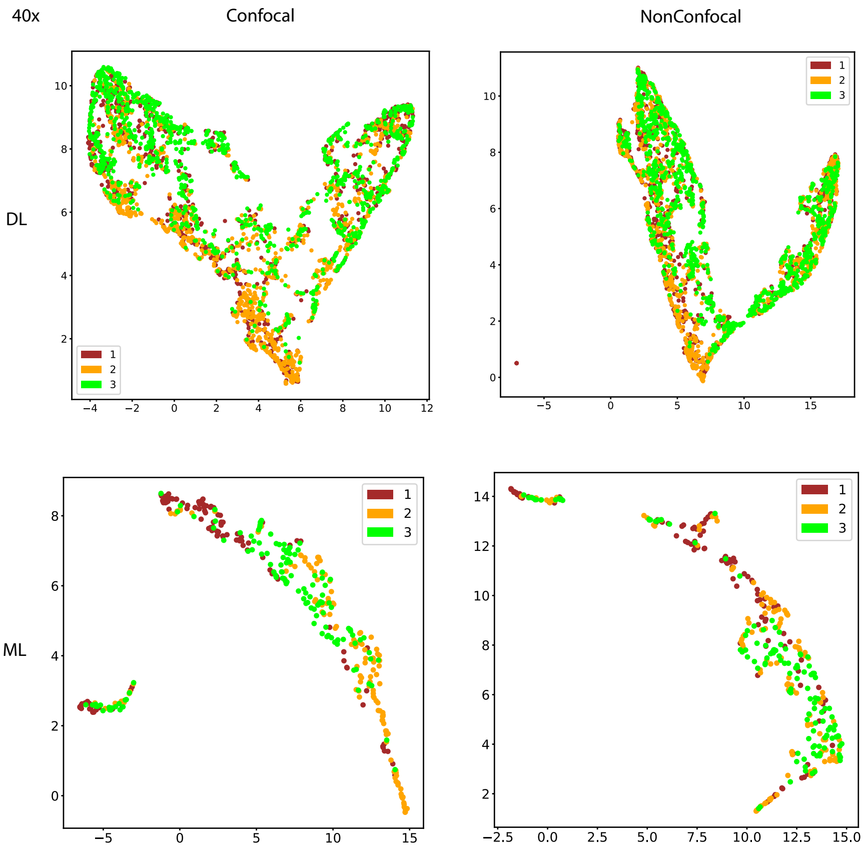


**Supplementary Figure 5: UMAP of embeddings of experiment 3 plates with confocal and non-confocal imaging.**
Every point corresponds to the embedding of an image. On the top using the CellDeathPred model which was trained on images from experiment 1 and 2. Individual wells are visualized as points on the scatter plot of the first two principal components.
On the bottom UMAP of 245 features extracted from the images initially extracted from Columbus software.
The color code is according to the plate category (red = "plate01", yellow = "plate02", green = "plate 03") and was added after the UMAP was conducted.
